# Prevalence and distribution of antimicrobial resistance determinants of *Escherichia coli* isolates obtained from meat in South Africa

**DOI:** 10.1101/626267

**Authors:** Ishmael Festus Jaja, James Oguttu, Ezekiel Green, Voster Muchenje

## Abstract

**Objective:** This study aimed to characterise antibiotics resistance of *Escherichia coli* isolates from the formal meat sector (FMS) and informal meat sectors (INMS).

**Method:** A total of 162 and 102 *E. coli* isolates from the FMS, and INMS respectively were isolated by standard culture-based, and biochemical reactions. The isolates were further confirmed by polymerase chain reaction (PCR). The disc diffusion method was used to screen for antimicrobial susceptibility against 19 different antibiotics. The presence of class 1-2 integrons in each *E. coli* isolates was assessed using 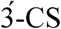 and 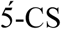 regions specific primers.

**Result:** Among the 19 antimicrobials, resistance to tetracyclines, aminoglycosides, cephalosporins, and nitrofurans were found to be more frequent than carbapenems and phenicol with a noticeable increase in the number of multi-drug resistance ranging from three to ten antimicrobials. A total of 20 resistance determinants were assessed with their prevalence and distributions obtained as follows for FMS and INMS respectively; [aminoglycosides: *aadA* (40.6%; 31.9%), *aacC2* (21.4%; 31%), *aphA1* (20.8%; 15.1%), *aphA2* (37.7%; 18.9%) and *strA* (6.5%; 9.4%)], [β-lactams: *ampC* (20%; 45%), *blaTEM*, (4.4%; 13.3), and *blaZ* (8.9%; 2.2%)], [Chloramphenicol: *catI* (1.7%; 1.7%), and *cmIA1* (1.7%; 1.7%)] and [tetracyclines: *tetA* (7.7%; 15.4%), *tetB* (11.5%; 24%), and *tetM*, (1.9%; 8.7%)], and [sulfonamides: sul1 (22.2%; 26.7%), sul2 (17.8%; 6.7%)].

**Conclusion:** Multiple antibiotic resistance (MAR) indexes ranged from 0.2 - 0.5. The results reveal a high prevalence of multidrug-resistant *E. coli* isolates and resistance determinants suggesting that consumers and handlers of such meat are at risk of contracting antibiotic resistant *E. coli*-related foodborne disease.

## 1 Background

Meat is a high-risk food, and many studies have shown a causal relationship between meat consumption and disease outbreaks. Meat-related foodborne diseases frequently occur due to the consumption of *Escherichia coli* contaminated raw or poorly processed meat. The United States, Center for Disease Control and Prevention (CDC) estimates for the year 2011 that one in six or forty-eight million people are infected with a foodborne illness each year, resulting in 3000 deaths. Of this number, *E. coli* O157: H7 caused an estimated 73480 illnesses each year, resulting in more than 2000 hospitalisations and 60 deaths [1].

Even though raw food hygiene and food safety epidemiology is still in its infancy in Africa, foodborne disease (FBD) is common in developing countries. For instance, 10200 cases of Shiga toxin *E*.*coli* (STEC) related FBD was reported in WHO subregion AFR D and E circa 2012 [2]. The actual prevalence of food-borne infections is difficult to determine, primarily because only a small percentage of incidence is officially reported. Even when cases of foodborne infections are reported, only in a limited number is the aetiology determined [3].

Antibiotics play a vital role in the treatment and management of bacterial infections, leading to a reduction in morbidity and mortality of both human and animal patients. However, the misuse of antibiotics in agriculture, veterinary and medical enterprises drives the selection of antibiotic-resistant bacteria that resist and overcome the action of the antibiotic. Approximately, 80% of all antibiotics used worldwide are in agriculture and aquaculture [4]. In livestock husbandry, antibiotics are used for prevention of infection or the simultaneous treatment of healthy and sick animals in a group during an outbreak of disease. It can further be used as antimicrobial feed additives (AFAs) for growth promotion and performance in production animals [4,5].

In 2006, the European Union (EU) prohibited the use of AFA for growth promotion in swine, cattle, poultry, and rabbits. Similarly, the US Food and Drug Administration opted for the voluntary phasing out of antibiotics for growth promotion (AGP) in animals in the USA in December 2013 [4,6]. However, in many developing countries, no such bans or plans to phase out AGPs have been put in place [4] and yet many of these countries bear the heavy burden of foodborne diseases caused by resistance pathogens.

Although epidemiological surveillance has established an association between the usage of antibiotic and antimicrobial resistance [7], South Africa registered sixty-four antimicrobial products, representing nineteen active pharmaceutical ingredients registered as in-feed mixes for growth promotion in the years 2002-2004. This included WHO-banned antimicrobials such as virginiamycin, spiramycin, tylosin, and bacitracin [7]. Meaning that South Africa still permits the use of these compounds as antibiotic feed additives, hence, it is no wonder that AFAs constituted two-thirds of all antimicrobials sold for animal use over these three years [4,5].

South Africa faces the challenge of scarcity of veterinary expertise needed to provide professional veterinary care, surveillance and monitoring of antimicrobial use by farmers. The shortage of veterinary surgeon in the country was one of the reasons the Fertilisers, Farm, Feeds, Agricultural Remedies and Stock Remedies Act (Act 36 of 1947) allows farmers to purchase and use some antimicrobial agents without needing to produce a veterinary prescription. As a result, farmers can purchase and administer many of the prescribed veterinary antibiotics since they can be obtained as over-the-counter (OTC) stock remedies [8]. It is then no surprise that in 2002-2004, 72% of antimicrobials used in animals in South Africa were permitted by the Stock Remedies Act.

*Escherichia coli* is a ubiquitous gut microorganism which forms part of the natural flora of the gastrointestinal system. Its ability to acquire both resistant determinants and virulence factors has been acknowledged by many researchers [9]. Some *E*.*coli* strains are highly pathogenic and are categorised based on their virulence into different pathogroups. These pathogroups include enterotoxigenic *E. coli* (ETEC); enteropathogenic *E. coli* (EPEC); Shiga toxin-producing *E. coli* (STEC), enteroaggregative *E. coli* (EAEC), diffusely adherent *E. coli* (DAEC); and enterohemorrhagic *E. coli* (EHEC); a sub-class of enteroinvasive *E. coli* (EIEC); neonatal meningitis *E. coli* (NMEC), enteroaggregative *E. coli* and uropathogenic *E. coli* (UPEC) [10]. The pathogenicity of each pathotype depends on its ability to adhere, colonise, and invade the host’s cells system causing cell death and apoptosis. The secretion and transportation of cell surface molecules, siderophore formation, and toxins are other methods of establishing virulence in the host [1].

Foodborne diseases (FBD) associated with resistant *E. coli* has reached an alarming proportion and remains a global public health crisis [11]. The transfer of resistance to enteric and commensal bacteria enhancing pathogenicity when consumption of contaminated food and water occur is even a bigger problem. The emerging resistance to WHO classified critically important antimicrobial such as carbapenems, extended-spectrum cephalosporins (ESCs), aminoglycosides and fluoroquinolones (FQs) among Enterobacteriaceae remains worrisome [12]. Studies’ comparing the AMR in the formal and informal meat sectors are limited. Hence, constant surveillance of the resistance profile of the bacteria is a useful early warning epidemiological indicator. This study aimed to determine the phenotypic and genotypic profile of antimicrobial resistant *E. coli* isolates obtained from abattoir and slaughter points the formal and informal meat sectors in the Eastern Cape Province of South Africa.

## 2 Material and methods

### 2.1 Sample site and collection

Samples from the formal meat sector (FMS) were collected at three high throughput abattoirs located in the East London (HT1), Queenstown (HT2) and Port Elizabeth (HT3) from the year 2015 to 2016. During the same period (2015-2016) samples were also collected from the informal meat sector (INMS) in different towns such as Alice town (AT), King Williams’ town (KWT), and Cala town (CT) (Figure 1). Carcasses from the informal sector were slaughtered for traditional use, home use, and the informal meat market. A total of 83 and 35 carcasses were sampled in the FMS and INMS respectively by swabbing the rump, neck, brisket, and flank areas. Sterile throat cotton swab moistened with peptone water was used to swab a 100cm^2^ carcass surface of beef, mutton, and pork. All samples were transported in a cooler box to the laboratory for the detection and confirmation of the presence of *E. coli*. In total, 332 and 140 samples were collected from the formal and informal meat sector respectively. A total of 162 and 102 *E. coli* isolates were confirmed by molecular method (PCR) in formal and informal meat sector respectively (Figure 2). All the confirmed (n = 162 + 102) confirmed isolates were stored in glycerol for further antimicrobial susceptibility testing.

**Figure 1:**
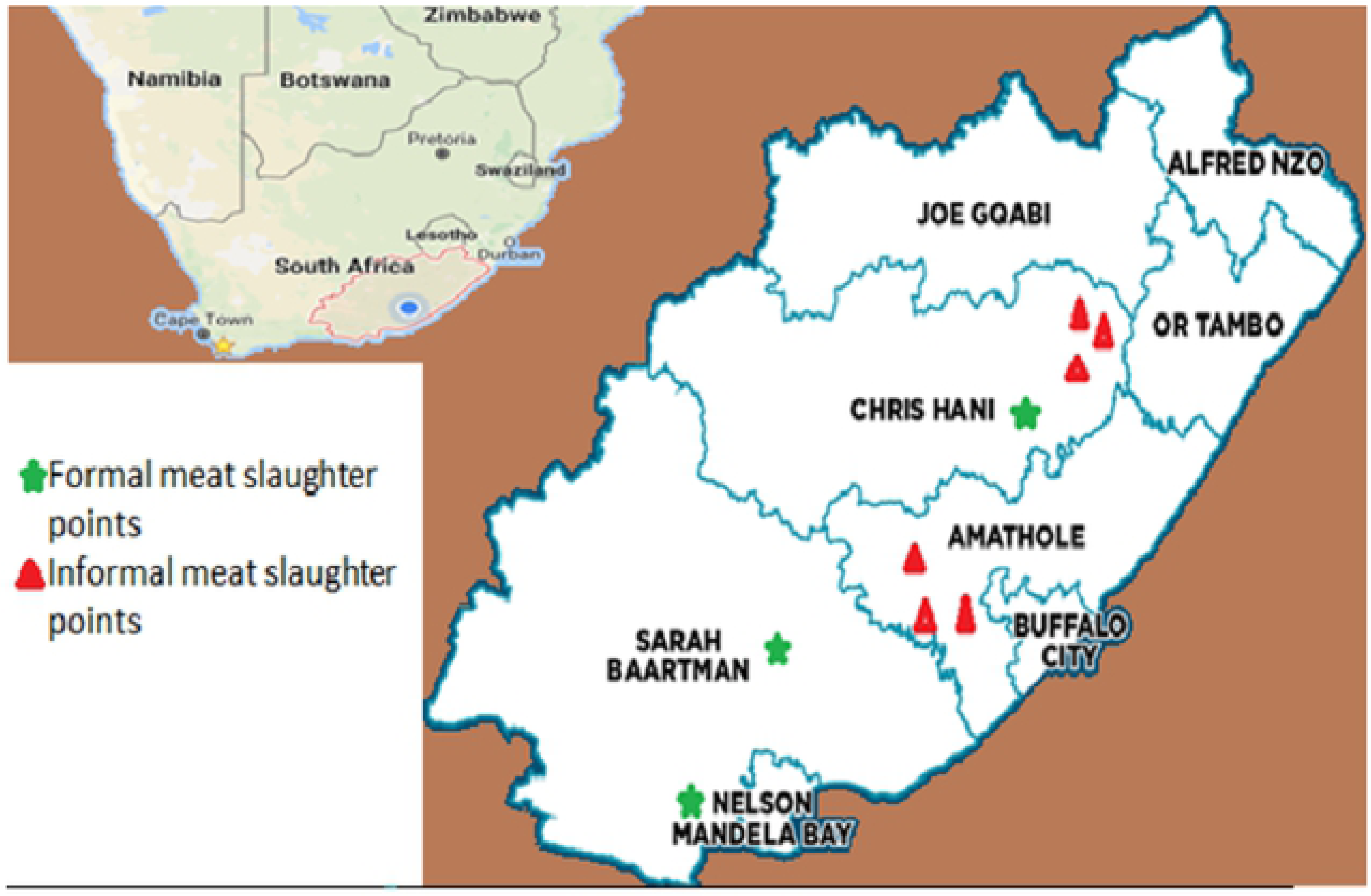
Map of the Eastern Cape Province showing sampling points.

**Figure 2:**
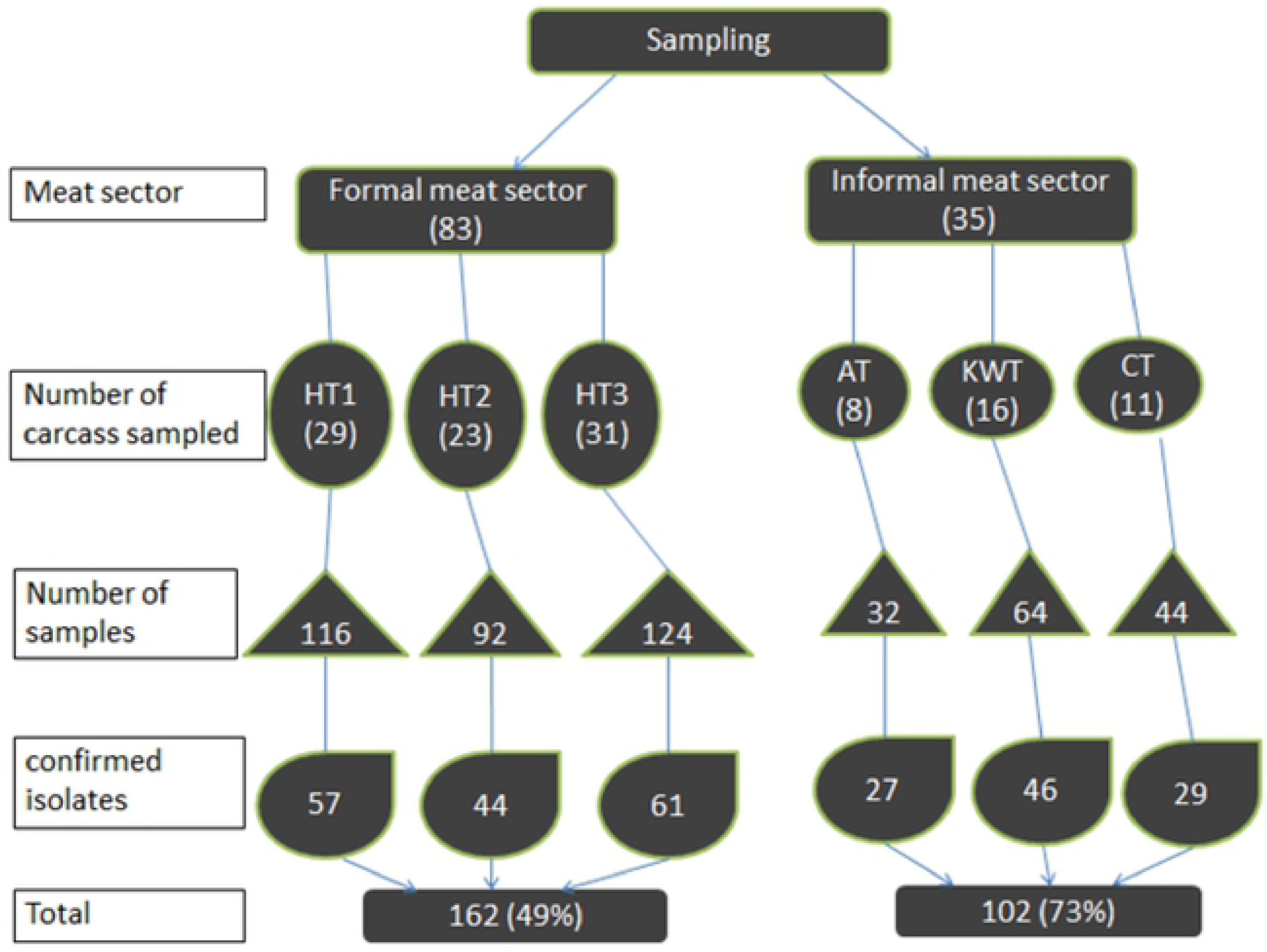
Flow diagram detailing the sample collection from the formal and informal meat sector.

### 2.2 Isolation and identification of *Escherichia coli*

All the swabs from all sampling sites were inoculated into tryptic soy broth (TSB) and were incubated for 24 hours at 37°C. Samples from the tryptic soy broth were then inoculated onto Eosin Methylene Blue agar (EMB) on different plates and incubated for 24-48 hours at 37°C. The EMB is a selective enrichment media and differential for *E. coli*. Single pure green metallic sheen colonies characteristic of *E. coli* on EMB were confirmed as presumptive isolates and stored in 30% glycerol awaiting further analysis.

### 2.3 DNA extraction

The presumptive *E. coli* isolates previously stored in glycerol were resuscitated and plated on EMB plates. Five distinctive colonies were picked from the EMB plates and inoculated into the nutrient broth and incubated for 48 hours at 37°C. DNA was extracted by the boiling method [13]. Briefly, 1 ml of broth solution containing *E. coli* was transferred to an Eppendorf tube and centrifuged (Thermo Fisher Scientific, Germany) for 15 mins at a speed of 13000 rpm. The supernatant was discarded, and the pellet was retained. Again another 1 ml of the broth was added and centrifuged at the same speed and duration; this was done five times to obtain a sizeable pellet. To wash the pellet; 200 µL of distilled water was added to the pellet and discarded. Again a volume of 200 µL of distilled water was added to the washed pellet, vortexed and centrifuged at a speed of 13000 rpm for 5 mins and the supernatant was discarded. The pellet was then placed on AccuBlock™ Digital Dry Baths (Labnet International, USA) at 100°C for 15 mins to lyse the cell. Cell debris was removed by centrifugation at 13000 rpm for 10 mins while the supernatant was stored as the DNA template.

### 2.4 PCR confirmation of *E. coli* isolates

Confirmation of presumptive *E. coli* isolates was by Polymerase chain reaction (PCR) in a total volume of 25µL containing 5.0µL of the DNA template, 5.5µL nuclease-free water, 12.5µL master mix, 1.0µL forward primer, and 1.0µL reverse primer. The *UidA* primers were used for PCR testing of the bacterial isolates. *Escherichia coli* ATCC 25922 served as the positive control strain [14]. The PCR condition is as follows: Initial denaturation at 94°C for 2 mins followed by 25 cycles of denaturation at 94°C for 1 min, annealing at 58°C for 1 min and extension at 72°C for 1 min. A final extension at 72°C for 2 mins. Holding was at 4°C (Table 1).

**Table 1:**
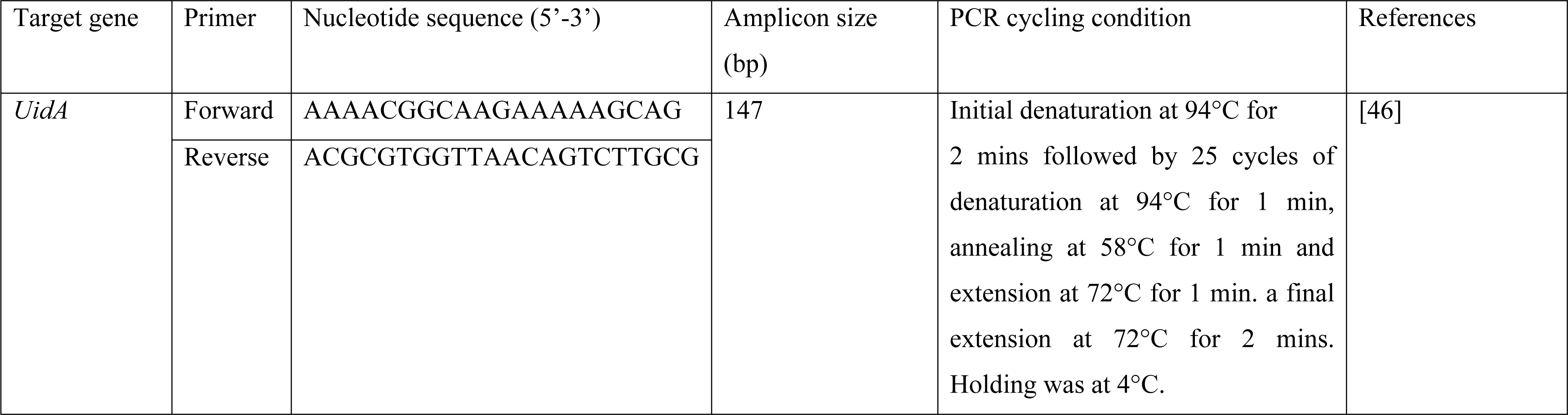
Primer sequence and PCR cycling condition of the targeted gene that confirms *E. coli*.

### 2.5 Antibiotic susceptibility profiles of *E. coli* isolates

The antimicrobial resistance profiles of confirmed isolates were determined using the Kirby– Bauer disk diffusion method on Mueller–Hinton agar [14]. Isolates were inoculated onto nutrient agar and incubated at 37 °C for 24 hours. A single colony was picked from the nutrient agar plate and suspended into 0.9 saline water and adjusted to give a reading of 0.5 McFarland turbidity standard. A 0.1 ml volume of the 0.5 McFarland suspension was swabbed evenly in at least three directions on the surface of a Mueller–Hinton agar plate. The surface of each plate was left to dry up, and then antimicrobial disks for each antimicrobial were placed at a specific place on the surface of the agar.

The plates were incubated lid side up at 37 °C for 24 hours. The zone of inhibition was recorded by measuring the size of the zone of inhibition around the disk. Isolates were classified as being resistant, intermediate and sensitive based on the Clinical and Laboratory Standards Institute guidelines [15]. Isolates which fell in the intermediate category were reclassified as being resistant [16].

Based on the CLSI guidelines, the following antibiotics were used for the antibiotic susceptibility test: Cotrimoxazole (25µg), Ciprofloxacin (5µg), Norfloxacin (10µg), Amoxicillin (30µg), Ampicillin (25µg), Tetracycline (30µg), Gentamicin (10µg), Streptomycin (300µg), Kanamycin (30µg), Neomycin (10µg), Ceftriaxone (30µg), Cefotaxime (30µg), Ceftazidime (10µg), Imipenem (10µg), Meropenem (10µg), Ertapenem (10µg), Doripenem (10µg), Chloramphenicol (30µg), Nitrofurantoin (300µg).

### 2.5 Detection of antimicrobial resistance genes

Specific primer sequences for the various resistance gene coding the phenotypic resistance of isolates observed were subjected to PCR assay as previously described [13]. Table 2 summarises the details of the process of gene sequencing. For cycling, a Bio-rad thermal cycler (Bio-Rad Mycycler, USA) was used. For antimicrobial classes such as sulphonamides, beta-lactams, tetracyclines, aminoglycosides, and phenicols, isolates were tested for the possession of various genotypic resistance determinants e.g. *aac(3)-IIa (aacC2), aph(3)-Ia (aphA1), aph(3)-IIa (aphA2), aph(3)-Ia (aphA1), aph(3)-IIa (aphA2), aadA, strA, blaTEM, blaZ, ampC, cat1, cat2, cmlA1, sul1, sul2, tetA, tetB, tetC, tetD*, and *tetM*.

**Table 2:**
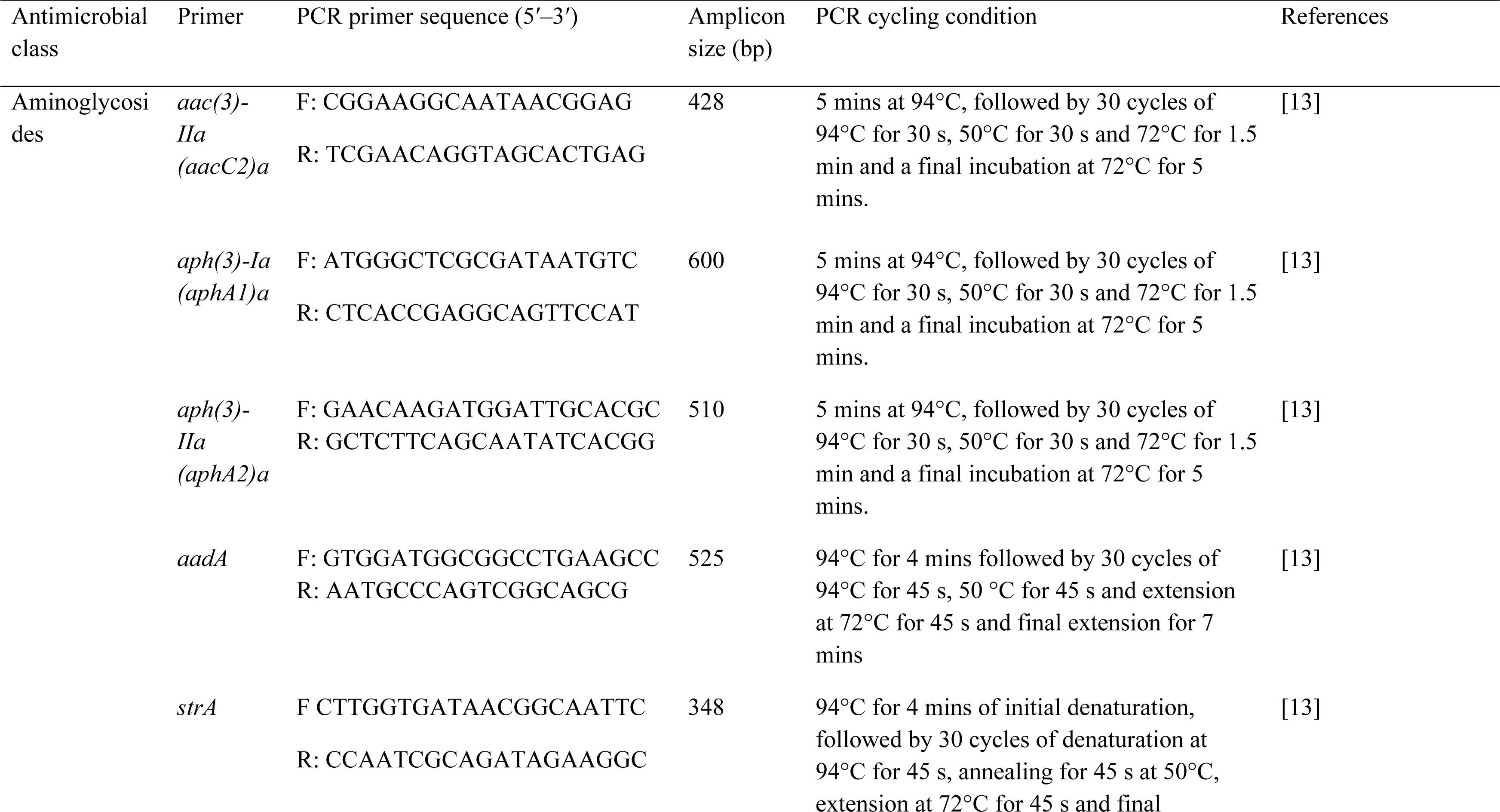
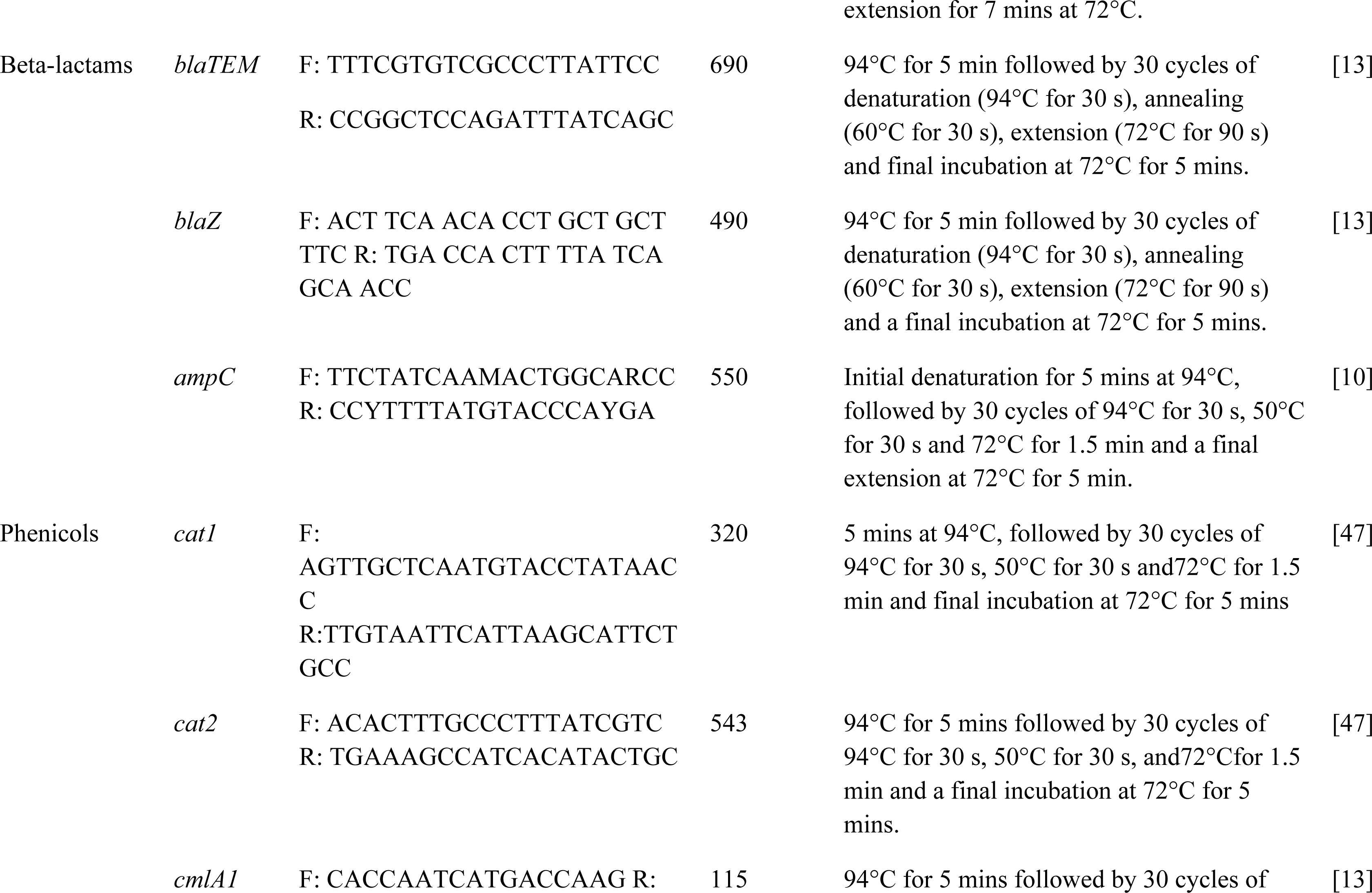
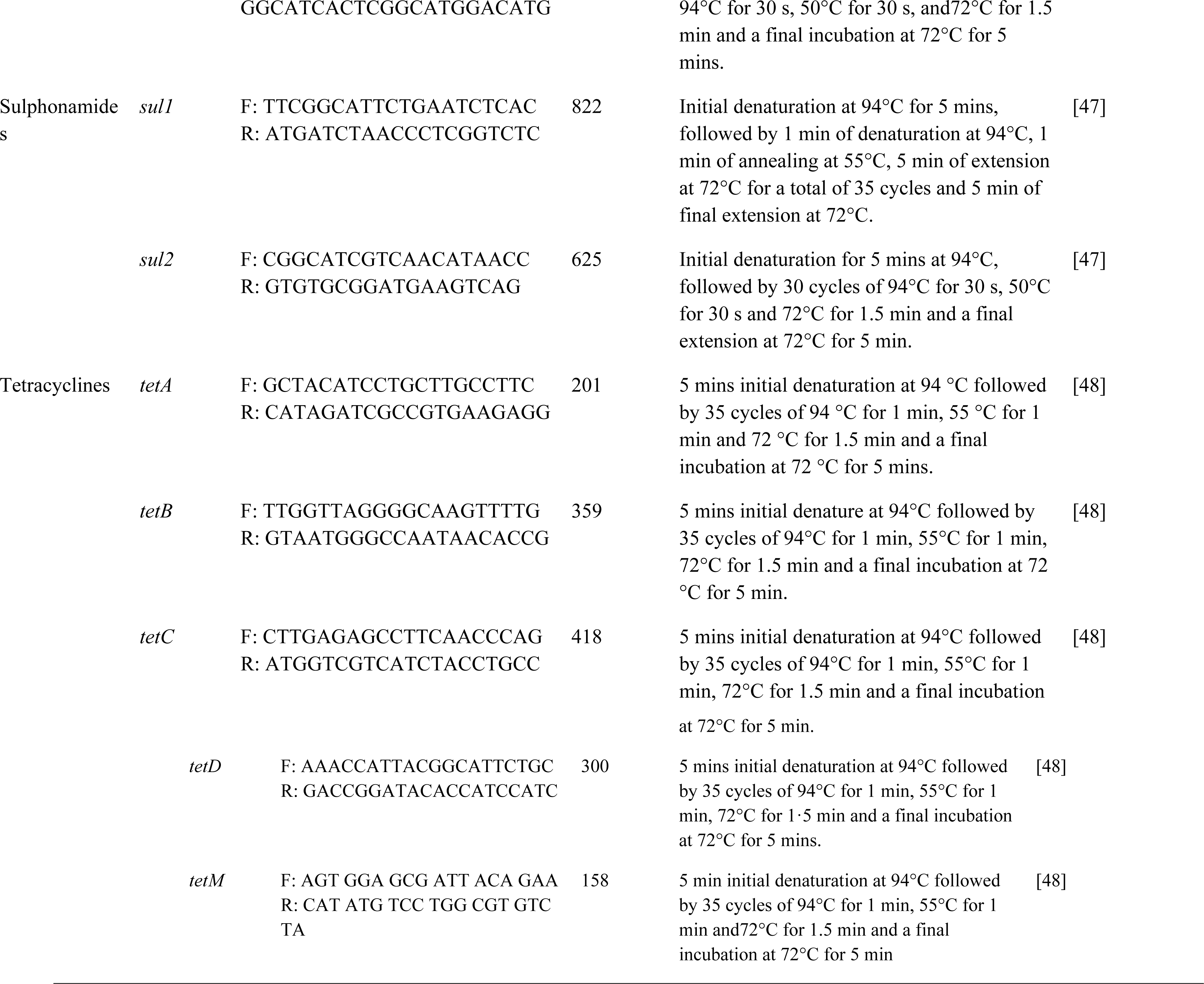
Primers set for antimicrobial resistance gene detection.

### 2.6 Gel electrophoresis

The amplified products were visualised by ethidium bromide staining after gel electrophoresis of 10 µL of the final reaction mixture in 1.5% agarose for 45 mins.

### 2.6 Statistical analysis

The data was captured in Microsoft Excel® (Microsoft Corporation, USA) and analysed using SPSS software (Version 24. IBM SPSS Inc, United States). The data were analysed to test for correlation between antibiotics resistance properties of *E. coli* isolate in the formal and informal meat sector. Statistical significance was set at P value < 0.05. Multiple antibiotics resistance index was calculated using the formula:

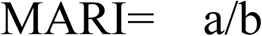

Where (a) is the aggregate antibiotic resistance score of all isolates from the sample, (b) is the total number of antibiotics used [17]. A MAR index ≥ 0.2 indicates the high-risk environment where antibiotics are often used [13].

## 3 Results

### 3.1 Isolation of *Escherichia coli*

From 83 carcasses that were sampled in the present study from formal meat sector, 322 samples were obtained. The 322 samples then yielded 162 confirmed *E. coli* isolates, of which 57, 44 and 61 isolates originated from HT1, HT2, and HT3 respectively (Figure 2). With regard to the informal meat sector, 35 carcasses were sampled, and these gave us 140 samples. The 140 samples then yielded 102 confirmed *E. coli* that included 27, 46, and 29 isolates from AT, KWT, and CT respectively.

### 3.2 Antimicrobial Susceptibility Testing

Antimicrobial resistance rate for *E. coli* isolates for the formal meat sector was as follows (Table 3): streptomycin 54.9% (89/162); ceftriaxone 54.9% (89/162); tetracycline 43.8% (71/162); nitrofurantoin 40.1% (65/162); neomycin 35.2% (57/162); amoxicillin 22.8% (37/162); ceftazidime and chloramphenicol 21.6% each (35/162); and kanamycin 20.4% (33/162). The proportion of drug resistance in isolates from the informal meat sector was as follows (Table 3): streptomycin 48.0% (49/102); tetracycline 32.4% (33/102); neomycin 30.4% (31/102); chloramphenicol 24.5% (25/102); gentamicin 22.5% (23/102); imipenem 21.6% (22/102); ceftriaxone 20.6% (21/102); kanamycin 19.6% (20/102); and cotrimoxazole 15.7% (16/102). Multiple antibiotic-resistant phenotypes (MARPs) pattern for isolates from the formal meat sector ranged from 1-5 (MARI, 0.2-0.5) and MARPs for the informal meat sector ranged from 2-15 (MARI, 0.2-0.5) (Table 4).

**Table 3:**
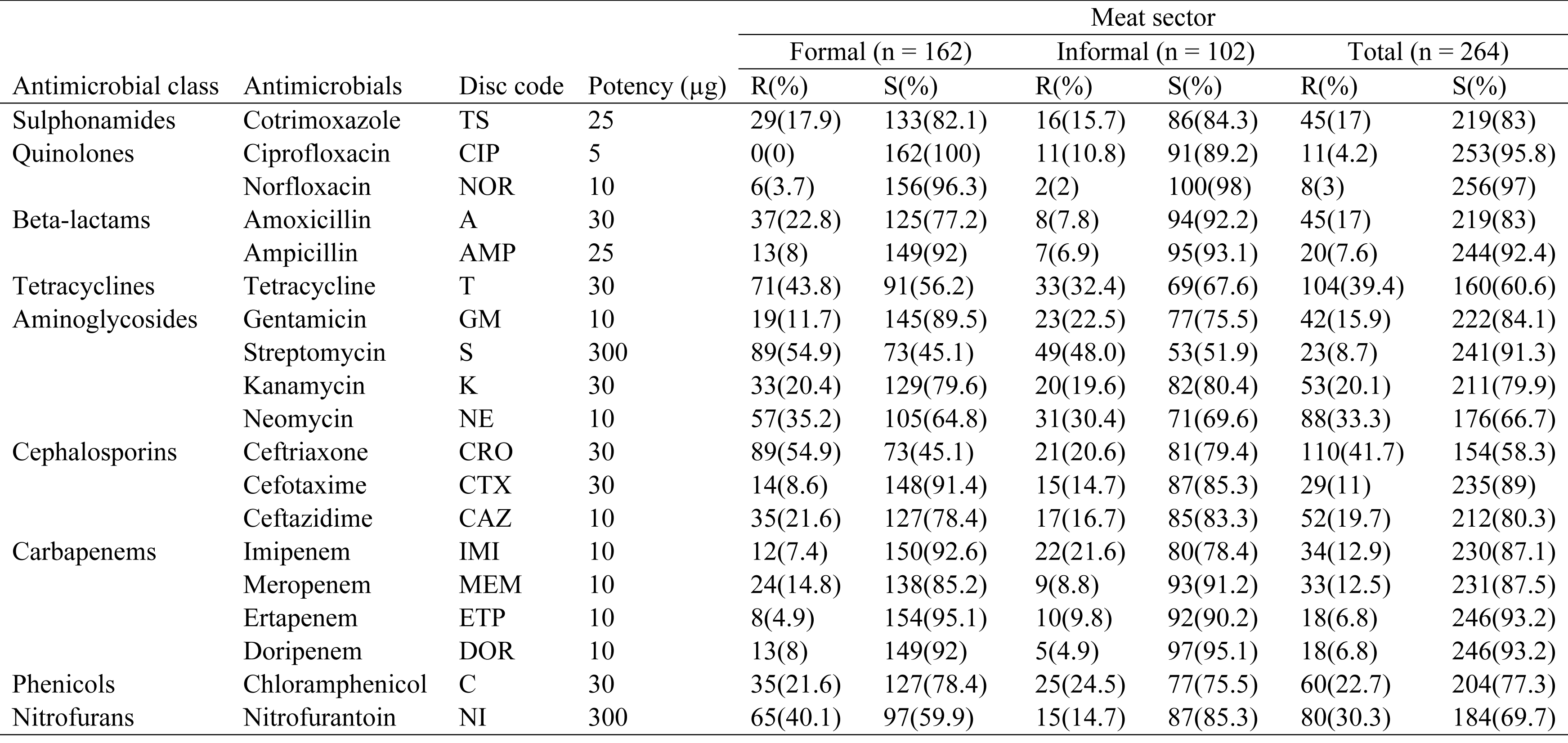
Antibiotic susceptibility pattern of *E. coli* isolates.

**Table 4:** Multiple antibiotic-resistant phenotypes (MARPs) pattern of *E. coli* isolates from the formal and informal meat sector

| Pattern number | Number of Antibiotics | Pattern | Meat sector |  | MARI |
| --- | --- | --- | --- | --- | --- |
|  |  |  | Formal | Informal |  |
| 1 | 3 | T-TS-CIP | 3 | 2 | 0.2 |
| 2 | 3 | A-AMP-CIP | 5 | 3 | 0.2 |
| 3 | 3 | TS-AMP-A | 1 | 0 | 0.2 |
| 4 | 3 | GM-T-C | 5 | 15 | 0.2 |
| 5 | 4 | GM-AMP-A-TS | 2 | 3 | 0.2 |
| 6 | 4 | S-AMP-NI-CAZ | 1 | 0 | 0.2 |
| 7 | 4 | CAZ-CTX-CRO-TS | 3 | 5 | 0.2 |
| 8 | 5 | ETP-MEM-IM-DOR-CRO | 2 | 6 | 0.3 |
| 9 | 5 | TS-AMP-A-C-CAZ | 0 | 3 | 0.3 |
| 10 | 6 | MEM-CAZ-DOR-NI-T-C | 1 | 2 | 0.3 |
| 11 | 6 | K-AMP-GM-A-T-NE | 2 | 0 | 0.3 |
| 12 | 6 | MEM-NI-S-T-A-TS | 2 | 2 | 0.3 |
| 13 | 7 | A-AMP-TS-GM-IMI-NI-C | 0 | 4 | 0.4 |
| 14 | 7 | K-GM-MEM-NE-T-A-NOR | 8 | 3 | 0.4 |
| 15 | 8 | ETP-C-IMI-T-A-AMP-CTX-S | 4 | 5 | 0.4 |
| 16 | 8 | AMP-T-TS-NI-GM-CRO-ETP-A | 0 | 2 | 0.4 |
| 17 | 8 | CRO-ETP-A-MEM-IMI-NI-AMP-T | 0 | 1 | 0.4 |
| 18 | 9 | T-NE-K-AMP-NOR-TS-S-ETP-CTX | 2 | 3 | 0.5 |
| 19 | 10 | NI-CIP-AMP-T-TS-C-S-GM-IMI-A | 0 | 2 | 0.5 |
| 20 | 10 | C-GM-TS-T-CRO-ETP-NOR-NI-A-AMP | 1 | 1 | 0.5 |

### 3.3 Antimicrobial resistance genes and pattern of resistance

Aminoglycoside resistance determinants were the most common (Table 5) and were as follows for isolates from the formal sector: streptomycin (*aadA*: 40.6%); kanamycin (*aphA1*: 20.8; *aphA2*: 37.7%); gentamycin (*aacC2*: 21.4%). Other resistant determinants that were common include the ones for cotrimoxazole (sul1: 22.2%) and (sul2: 17.8); ampicillin (*ampC*: 20%); and tetracycline (*tetB*: 11.5%).

**Table 5:** Percentage and distributions of antimicrobial resistance determinants among *E. coli* from formal and informal sector

| Antimicrobial agent | Disc code | Antimicrobial resistance gene | Gene group and or general function | Formal (%) | Informal (%) | Total (%) |
| --- | --- | --- | --- | --- | --- | --- |
| Gentamicin (n = 42) | GM | *aac(3)-IIa (aacC2)a* | Aminoglycoside resistance | 9(21.4) | 13(31) | 22(52.4) |
| Kanamycin (n = 53) | K | *aph(3)-Ia (aphA1)a* | Aminoglycoside resistance | 11(20.8) | 8(15.1) | 19(35.8) |
|  |  | *aph(3)-IIa (aphA2)a* | Aminoglycoside resistance | 20(37.7) | 10(18.9) | 30(56.6) |
| Neomycin (n = 88) | NE | *aph(3)-Ia (aphA1)a* | Aminoglycoside resistance | 8(9.1) | 13(14.8) | 21(23.9) |
|  |  | *aph(3)-IIa (aphA2)a* | Aminoglycoside resistance | 20(22.7) | 10(11.4) | 30(34.1) |
| Streptomycin (n = 138) | S | *aadA* | Aminoglycoside resistance | 56(40.6) | 44(31.9) | 100(72.5) |
|  |  | *strA* | Aminoglycoside resistance | 9(6.5) | 13(9.4) | 22(15.9) |
| Amoxicillin (n = 45) | A | *blaTEM* | Beta-lactam | 2(4.4) | 6(13.3) | 8(17.8) |
|  |  | *blaZ* | Beta-lactam | 4(8.9) | 1(2.2) | 5(11.1) |
| Ampicillin (n = 20) | AMP | *ampC* | Beta-lactam - *AmpC* | 4(20) | 9(45) | 13(65) |
| Chloramphenicol (n = 60) | C | *catI* | Chloramphenicol resistance | 1(1.7) | 1(1.7) | 2(3.3) |
|  |  | *cat2* | Chloramphenicol resistance | 1(1.7) | 0(0) | 1(1.7) |
|  |  | *cmlA1* | Efflux pump | 1(1.7) | 1(1.7) | 2(3.3) |
| Cotrimoxazole (n = 45) | TS | *sulI* | Sulphonamide resistance | 10(22.2) | 12(26.7) | 22(48.9) |
|  |  | *sul2* | Sulphonamide resistance | 8(17.8) | 3(6.7) | 11(24.4) |
| Tetracycline (n = 104) | T | *tetA* | Tetracycline resistance | 8(7.7) | 16(15.4) | 24(23.1) |
|  |  | *tetB* | Tetracycline resistance | 12(11.5) | 25(24) | 37(35.6) |
|  |  | *tetC* | Tetracycline resistance | 1(1) | 7(6.7) | 8(7.7) |
|  |  | *tetD* | Tetracycline resistance | 1(1) | 13(12.5) | 14(13.5) |
|  |  | *tetM* | Tetracycline resistance | 2(1.9) | 9(8.7) | 11(10.6) |

Resistance determinant found in the informal sector was streptomycin (*aadA*: 31.9%); kanamycin (*aphA1*: 15.1%) and (*aphA2*: 18.9%); neomycin (*aphA1*: 14.8%); and gentamycin (*aacC2*: 31%). Others were cotrimoxazole (sul1: 26.7%); amoxicillin (*blaTEM*: 13.3%); ampicillin (*ampC*: 45%); and tetracycline (*tetA*: 15.4%; *tetB*: 24%; *tetD*: 12.5%).

The number of genotype resistance determinants pattern ranged from 2-4 for the various antibiotic genes tested. In the formal sector, 1-4 isolates were found to have multiple genes coding for resistance determinants. However, in the informal sector, 1-12 isolates possessed multiple genetic components for antimicrobial resistance determination (Table 6).

**Table 6:** Genotypic resistance determinants profile of *E. coli* isolates from the formal and informal meat sector

| Antimicrobial class | Number of pattern | Genotype resistance determinants pattern | Formal | Informal | Total |
| --- | --- | --- | --- | --- | --- |
| Aminoglycosides | 2 | *aac(3)-IIa (aacC2)<sup>a</sup>+aph(3)-Ia (aphA1)<sup>a</sup>* | 1 | 1 | 2 |
|  | 3 | *aac(3)-IIa (aacC2)<sup>a</sup>+aph(3)-Ia (aphA1)<sup>a</sup>+aph(3)-IIa (aphA2)<sup>a</sup>* | 2 | 0 | 2 |
|  | 2 | *aadA+strA* | 4 | 1 | 5 |
| Beta-lactams | 2 | *blaTEM+blaZ* | 0 | 1 | 1 |
|  | 2 | *blaZ+ampC* | 0 | 1 | 1 |
| Phenicol | 2 | *catI+catII* | 1 | 0 | 1 |
|  | 2 | *catII+cmlA1* | 1 | 0 | 1 |
| Sulphonamides | 2 | *sulI+sul2* | 2 | 6 | 8 |
| Tetracyclines | 2 | *tetA+tetB* | 1 | 4 | 5 |
|  | 2 | *tetA+tetC* | 3 | 9 | 12 |
|  | 2 | *tetB+tetD* | 0 | 1 | 1 |
|  | 3 | *tetA+tetB+tetD* | 0 | 5 | 5 |
|  | 3 | *tetC+tetA+tetB* | 2 | 4 | 6 |
|  | 3 | *tetA+tetB+tetM* | 1 | 2 | 3 |
|  | 3 | *tetB+tetD+tetM* | 2 | 0 | 2 |
|  | 3 | *tetD+tetC+tetA* | 1 | 0 | 1 |
|  | 4 | *tetA+tetB+tetC+tetD* | 1 | 4 | 5 |
|  | 4 | *tetA+tetB+tetC+tetM* | 1 | 3 | 4 |
|  | 4 | *tetB+tetC+tetD+tetM* | 2 | 4 | 6 |

<sup>a</sup> Alternative nomenclatures are in parentheses

## 4 Discussion

Livestock is recognised as a primary reservoir of various pathotypes of *Escherichia coli*, which have been epidemiologically linked to many incidences of meat-related food-borne diseases [18]. Hence, monitoring the microbial quality of meat using indicator organisms such as *E. coli* is of medical and veterinary importance [19]. The development of antimicrobial resistance among pathogens that impact human and animal health further buttresses the need for intensified surveillance. Thus *E. coli* due to its existence in the gastrointestinal tract of animals and its ability to acquire antimicrobial resistance has been designated as a sentinel organism in antimicrobial resistance surveillance programs worldwide [20].

In the current study, meat processed in the formal (49%) and informal (73%) harboured *E. coli* (Figure 2). These results suggest that contamination of meat does occur during slaughter, and thus highlights the quality of meat processed in both the formal and informal meat sectors. The result of this study agrees with the 74%, 57.9% and 43% prevalence of *E. coli* found from beef in South Africa, Colombia and Burkina Faso respectively [21–23]. However, a low prevalence of 1-4% and 2.8% of *E. coli* was reported in the city of Graz, Saudi Arabia and Amatole district of the Eastern Cape Province, South Africa respectively [24,25]. Both studies mainly focused on *E. coli* O157: H7, hence the differences in both studies could easily be explained given the very low levels of prevalence of *E*.*coli* observed compared to what we observed in the present study.

The authors expected meat from the formal sector to be of high microbial quality. However, results reported here indicated that just under half (49%) of the carcasses from the formal sector were positive for *E. coli*. Even though this was lower than the 73% from the informal market, it can be considered to be very high. Contamination of carcass at abattoirs could be a direct consequence of dirty hides or airborne pathogens, contaminated knife, aprons hands and other cutting tools [26]. On the other hand, animal slaughter in the informal sector is performed in an unhygienic, unorganised and unregulated environment. This explains why a higher level of contamination was observed among carcasses from the informal sector compared to those from the formal sector. In one study of meat safety knowledge among butchers during traditional meat slaughter, many meat handlers during traditional meat slaughter demonstrated moderate knowledge of meat safety rules [27]. However, in practice this is hardly true given that meat from the informal market is commonly associated with a greater risk of food poisoning [28].

Globally, the bacteriological safety of meat is a pressing public health concern. Slaughter and dressing with personnel practices carry the highest value when analysing the risk of contaminating the carcass. Thus a breach in hygiene protocol could lead to the introduction of microorganisms at the production, processing, and consumption of meat and meat products [27], hence the need for a risk-based food safety system.

In the formal meat (FMS), profoundly high phenotypic resistance was observed for streptomycin (54.9%), ceftriaxone (54.9%), tetracycline (43.8%) and nitrofurantoin (40.1%), all of which are antimicrobials commonly used in human medicine. On the other hand, antimicrobial resistance in the informal meat (INMS) sector tended to be highest for streptomycin (48.0%), neomycin (30.4), tetracycline (32.4%), chloramphenicol (24.5%), imipenem (21.6%) and ceftriaxone (20.6%). Although studies’ comparing the AMR in the formal and informal meat sector are limited, results of the present study are consistent with available studies on AMR in the meat sector that reported a sustained prevalence of AMR to streptomycin and tetracycline as well as the emerging AMR to cephalosporins [19].

The possible reason for the variability in the proportions of AMR in the FMS and INMS could be sample size, hygiene management systems, and the application of antibiotics for prophylaxis and metaphylaxis in livestock farms in both meat sectors. The other reason could be the co-selection of resistant determinant and co-resistance of the *E. coli* isolates. The simultaneous resistance to penicillin, streptomycin, tetracycline, erythromycin, kanamycin, and virginiamycin has been reported in studies conducted elsewhere [29–31]. A similar co-selection of sulphonamide resistance genes was reported in chickens treated with streptomycin [32]. Moreso, a Canadian study found a predominant pattern of AMR with extended-spectrum cephalosporin (ESC)-resistant *E. coli* strains, with co-resistance to streptomycin, cefoxitin, trimethoprim-sulfamethoxazole, sulfisoxazole, ampicillin, amoxicillin/clavulanic acid, chloramphenicol, and tetracycline [30]. Also, in an extensive study of *E. coli* isolates from various countries authors observed that nearly 75% of ampicillin-resistant *E. coli* isolates were also resistant to streptomycin and tetracycline [33]. Hence, the authors of that study suggested that the resistance genes for these drugs are linked on plasmids.

The prevalence of streptomycin resistance in this study could be linked to its extensive use for the treatment of bacterial infections of plants and animals [34,35]. Generally, aminoglycosides resistance is mediated by aminoglycoside-modifying enzymes, including acetyltransferases and nucleotidyltransferases, aminoglycoside phosphotransferases, and 16S rRNA methylases, all of which have been reported in Enterobacteriaceae [36]. The increasing prevalence of resistance against streptomycin has led to its designation as a critical epidemiological marker to indicate the likelihood of multidrug-resistance (MDR) in pathogens [34]. Streptomycin resistance is frequently mediated by *aadA* genes, which are typically present on integrons causing streptomycin adenylation [34,37]. Thus, it is not surprising that the *aadA* gene and to a lesser extent the *strA* gene were the main genetic element of resistance observed in the present study (Table 6.5).

Cephalosporins such as ceftriaxone, cefotaxime, and ceftazidime are used in food-producing animals, and this imposes greater selection pressure for the development of extended-spectrum β-lactamases (ESBL)-producing and multiple-antimicrobial-resistant *E. coli*. ESBLs are the main contributors to extended-spectrum cephalosporin (ESC) resistance in *E. coli* and transfer resistance to cephalosporins with an oxyimino side chain [38]. Resistance to extended spectrum cephaloporinases (ESCs) in *E. coli* has been linked with the extended-spectrum β-lactamases (ESBLs) and plasmid-mediated Ambler class C cephamycinases [39]. Cephalosporin resistance has also been associated with extensive use of antibiotics in clinical practice. It is highly possible that apart from cross-contamination of the carcass with animal faeces during slaughter, that many *E. coli* isolates in the INMS could originate from humans. Therefore ceftriaxone resistance in FMS and INMS could be due to the inherent danger of the spread of mobile genetic element of resistance through horizontal gene transfer.

In South Africa, Stock Remedies Act, 1947 permits tetracycline to be purchased over the counter (OTC) without a veterinary prescription [40]. Therefore tetracycline resistance in FMS and INMS could be a direct consequence of its widespread use in the treatment of bacterial infection and tick-borne diseases in livestock [5]. Ticks are endemic in the province where the study was conducted. Hence farmers routinely treat their animal with tetracycline prophylactically to prevent disease outbreak. Besides, low doses of tetracycline have been used to control weaning diarrhoea and also included in the feed as antibiotic feed additives (AFAs) for growth promotion [5]. The results observed in this study are in agreement with studies done in Ethiopia, Iran, and Pakistan [41–44]. Meanwhile, our findings contradict findings of earlier studies done in South Africa and Poland that reported lower prevalences [25,29].

Although the *tet A, B, C, D* and *M* genes were observed in both FMS and INMS, the burden of resistance was more in the INMS. The high prevalence of *tet* in INMS suggests the therapeutic overuse or misuse of tetracyclines by communal farmers who happen to be the are the main suppliers of animals slaughtered on the INMS. The result of the study were expected, considering that over 70% of antibiotics used in livestock production in South Africa can be purchased over the counter [4].

Although the use of chloramphenicol in veterinary medicine and aquaculture has been banned worldwide, ampicillin, cotrimoxazole and other antimicrobials are still commonly used in livestock production in South Africa for treatment, prophylaxis and growth promotion purposes [4,5]. The injudicious use of these drugs may exert selective pressure sustaining the emergence of resistant bacterial strains. Furthermore, co-selection of multiple resistance mechanisms through the use of various antibiotics is possible because resistance genes for many antimicrobial agents are placed on single conjugative plasmids [45]. Apart from the previously mentioned mechanism of multidrug resistance (MDR), inactivation or enzymatic degradation of antimicrobials and chemical transformation of antimicrobial compounds by glycosylation, adenylation, acetylation, phosphorylation, and hydroxylation have also become steadily more apparent as causes of MDR [37]. Some of these mechanisms might be responsible for resistance observed among pathogens studied in this study, and could also be responsible for the resistance to nitrofurantoin which is not commonly used in veterinary medicine in South Africa.

## 5 Conclusion

This study revealed a high burden of resistance against important antimicrobials such as streptomycin, neomycin, ceftriaxone, chloramphenicol, and tetracycline including imipenem and meropenem. Genes encoding cephalosporin resistance are commonly situated on self-transmissible plasmids which may be promiscuous and capable of disseminating into a broad range of microbiota. Furthermore, resistance of *E. coli* isolates to antibiotics of choice in human therapy such aminoglycoside, phenicols, carbapenems, and cephalosporins pose a grave danger for success in human chemotherapy. Resistance in nitrofurantoin, a drug not commonly in use in South Africa, suggests that factors other than selective pressure must have an impact on the emergence of resistant *E. coli*.

In this study, only *blaTEM* and *blaZ* genes were tested, but the possibility of the *E. coli* isolates harbouring other ESBL genes such as *blaCTX*, and *blaSHV* is highly plausible; suggesting that these isolates could potentially be dangerous to public health. Such risk becomes pronounced in compromised food systems as bacteria strains can be transferred to humans via the food chain. Aside the mobilisation of plasmids that can lead to transfer of resistance genes to other Gram-negative and commensal bacteria in the environment, the public health and veterinary concern regarding AMR warrants a sustained and concerted local, regional and international coordinated surveillance and containment system for effective prevention of AMR in food animals. Simple intervention strategies, such as the prudent use of antimicrobials, promoting regular intermittent washing of hand and knife with hot water and soap by slaughterhouse workers and good hygienic practices at the abattoirs and informal slaughterhouses, can have a profound impact on public health.

## Abbreviations

DAEC: diffusely adherent E. coli
EHEC: enterohemorrhagic E. coli
INMS: Informal meat sector
UPEC: uropathogenic E. coli
AFA: Antimicrobial feed additives
AGP: Antimicrobial growth promoters
AMR: Antimicrobial resistance
AT: Alice town
CDC: Center for Disease Control and Prevention
CT: Cala Town
EAEC: enteroaggregative E. coli
EAEC: enteroaggregative E. coli
EIEC: enteroinvasive E. coli
EMB: Eosin methylene blue
EPEC: enteropathogenic E. coli
ESCs: extended-spectrum cephalosporins
ETEC: enterotoxigenic E. coli
EU: European Union
FBD: Foodborne disease
FMS: Formal meat sector
FQs: fluoroquinolones
(HT1): High throughput abattoirs in the East London,
(HT2): Queenstown and
(HT3): Port Elizabeth
KWT: King William town
MARI: Multiple antimicrobial resistance index
MARPs: Multiple antimicrobial resistance phenotypes
NMEC: neonatal meningitis E. coli
STEC: Shiga toxin-producing E. coli
TSB: Tryptic soy broth
WHO: World Health Organisation

## Ethics approval and consent to participate

Ethical approval number MUC351SJAJ01 was obtained from the University of Fort Hare research ethics committee.

## Consent for publication

Written consent for publication was obtained from each participating abattoir prior to the microbial survey.

## Availability of data and material

The data that support the findings of this study are available from University of Fort Hare repositories. Data are however available from the authors upon reasonable request and with permission of the University of Fort Hare

## Authors’ contributions

IFJ designed and carried out the study, JO edited the manuscript and made useful technical and specialist inputs, EG and VM supervised the study.

## Acknowledgements

The authors would like to thank the participating abattoirs for approving the study and the South African National Research Foundation (NRF) for funding the project.

